# Sympathetic neurons secrete retrogradely transported TrkA on extracellular vesicles

**DOI:** 10.1101/2022.05.24.493096

**Authors:** Ashley J Mason, Austin B Keeler, Farah Kabir, Bettina Winckler, Christopher Deppmann

**Author notes:** Corresponding authors and lead contact: Bettina Winckler, Christopher Deppmann.

## Abstract

Neuronal derived extracellular vesicles (EVs) have been well described in the central nervous system; however, studies in the peripheral nervous system have largely focused on EVs derived from supporting cell types such as endothelial cells or glia. Here we isolate EVs derived from sympathetic neurons and characterize them using immunoblot assays, nanoparticle tracking analysis and cryo-electron microscopy. Sizing of sympathetic EVs reveal a predominant peak between 45-75 nm as well as a range of larger sizes (90 nm to >350 nm), possibly due to multiple biogenic origins. We identified TrkA, a receptor for nerve growth factor (NGF), as a cargo for sympathetic EVs. Furthermore, TrkA on EVs was phosphorylated, indicating activated TrkA receptor. TrkA binds NGF at the axonal tip and is endocytosed and transported to the soma in signaling endosomes. We therefore examined if TrkA originating in the axon tip was subsequently able to be packaged into EVs and secreted by the somatodendritic domain of neurons. Using a compartmentalized culture system, we found that TrkA derived from endosomes originating in the distal axon can be detected on EVs secreted from the somatodendritic domain. In addition, inhibition of classic TrkA downstream pathways, specifically in somatodendritic compartments greatly decreases TrkA packaging into EVs. Our results suggest a novel trafficking route for TrkA: it can travel long distances to the cell body, be packaged into EVs and secreted. Secretion of TrkA via EVs appears to be regulated by its own downstream effector cascades, raising intriguing future questions about novel functionalities associated with TrkA^positive^ EVs.

## Introduction

Extracellular vesicles (EVs) are small, secreted lipid bilayer-enclosed vesicles implicated in a variety of functions, ranging from cargo transport, to intercellular signaling^1,2,3,4^. The term “extracellular vesicle” encompasses a wide range of vesicles secreted from cells including apoptotic bodies, ectosomes, microvesicles, exosomes and exomeres^4,5^. EVs are derived from two main sources in the cell: the plasma membrane or the endolysosomal system. EVs that bud off from the plasma membrane are generally termed microvesicles or ectosomes and contain surface cargos that are enriched on the plasma membrane^6^. EVs derived from the endolysosomal system are generated when a multivesicular body, an endocytic organelle, fuses with the plasma membrane. Upon fusion, the intraluminal vesicles (ILV) contained within the MVB get released into the extracellular milieu and are known as exosomes^7,8^.

Although the study of EVs has greatly expanded over the past decade, very little is known about EVs secreted by peripheral neurons^9^. The majority of the research has focused on peripheral nerve regeneration where EVs derived from non-neuronal sources (i.e., macrophages, Schwann cells, or endothelial cells) influence axonal repair^10^. Only one study has shown that sympathetic neurons release EVs in response to KCl-induced depolarization^11^. Furthermore, peripheral neurons can contain axons that extend many microns out from the cell body. Whether cargoes that originate in these distal axons can be trafficked back to the cell body and secreted as EVs remains unknown. In addition, specific cargos being secreted via EVs from sympathetic neurons are largely not yet identified.

In this study we characterize EVs secreted by sympathetic neurons that are derived from mouse superior cervical ganglia. In accordance with the guidelines set forth by the International Society for Extracellular Vesicles (ISEV) in their position paper “Minimal Information for Studies of Extracellular Vesicles” (MISEV)^12,13^, we characterize the size and concentration of EVs secreted by sympathetic neurons using Western blot, cryo-electron microscopy and nanoparticle tracking analysis (NTA). Furthermore, using microfluidic devices and the neurocircuit tracer, WGA, we show that cargo originating in the distal axon can undergo retrograde transport to the soma where it is packaged and released in EVs.

The neurotrophin receptor, TrkA, binds to nerve growth factor (NGF) at its axon terminals, internalizes into a signaling endosome (SE), and is retrogradely trafficked back to the somatodendritic compartment to promote survival, synapse formation, and several other trophic functions^14–16^. Much effort by our group and several others has gone into determining the fate of the retrogradely transported TrkA SEs. Interestingly, it has been shown that the SE arrives in the soma as a multivesicular body (MVB)^17^. Morphologically, MVBs are defined by intraluminal vesicles (ILVs) that are formed through budding-in events from the endosomal limiting membrane. Functionally, MVBs are a mature, late endosomal compartment largely destined for degradation. However, it has become increasingly clear that an alternative fate is for MVBs to fuse with the plasma membrane to release their internal ILVs as exosomes. The fate of an MVB-residing receptor after fusion with the plasma membrane depends on whether it remains on the limiting membrane or is internalized into ILVs. Receptors on the limiting membrane will be incorporated into the plasma membrane after MVB fusion. Alternatively, receptors residing on ILVs will be secreted on exosomes after MVB fusion. Our previous and current work show that TrkA undergoes both fates. We have shown that the activated retrogradely trafficked TrkA SE can induce association with the cytoskeletal protein, Coronin-1a, which slows degradation of TrkA by evading fusion with the lysosome^18^. The TrkA SE instead undergoes Coronin-1a dependent recycling to the plasma membrane and subsequent re-internalization^18^ of TrkA, a pathway we have termed “retrograde transcytosis”. In this work, we show that the activated TrkA receptor internalized in the axon is secreted as EVs from the somatodendritic domain of sympathetic neurons, demonstrating a new trafficking route for TrkA. Furthermore, TrkA secretion on EVs is influenced by signaling downstream of TrkA activation.

## Results

### Sympathetic neurons release extracellular vesicles

P3 mouse sympathetic ganglia were dissociated and 100,000 to 150,000 cells were plated in a 12 well plate. Extracellular Vesicles (EVs) were isolated from the conditioned media (CM) of mouse sympathetic cultures after 7 days *in vitro* (7 DIV) using differential centrifugation (Fig. 1A)^13^. Because serum is known to contain EVs, we grew superior cervical ganglia (SCG) cells in serum-free media supplemented by Prime XV IS-21. After the initial low-speed centrifugation steps, the supernatant was transferred and spun at 20,000 x g to obtain the pellet, “P20”. The supernatant from the 20,000 x g spin was then spun at 100,000 x g to yield a pellet “P100”. The P20 fraction is usually quite heterogeneous, containing dense vesicles such as apoptotic bodies, large ectosomes/microvesicles, and aggregates of smaller vesicles^13^. The supernatant of the P20 fraction contains smaller, less dense vesicles and particles (e.g. exosomes), which are pelleted in the P100 fraction (Fig. 1A). Both the P20 and P100 fractions were resuspended in PBS for subsequent nanoparticle tracking analysis (NTA) (Fig. 1A,B). Quantification of NTA tracked particles showed a greater concentration of particles in the P20 fraction compared to the P100 fraction (Fig. 1C). Size distribution histograms from NTA show a mean diameter of 134 nm and 136 nm for the P20 and P100 fraction, respectively (Fig. 1 E,F). To support the notion that these fractions are enriched for EVs, we blotted against canonical EV markers: CD63, CD81, and Alix (Fig. 1D, Supplementary Fig. S1). All three EV-associated markers were detected in the cell pellet, P20 and P100 fractions of three independent mouse litters (litter L1-L3; Fig. 1D). Importantly, cytochrome C, a mitochondrial marker, and calreticulin, an ER resident protein, were not detected in the P20 and P100 fractions, indicating undetectable contamination by intracellular organelles (Fig. 1D, Supplementary Fig. S1). Lastly, neither CD63, CD81, Alix, calreticulin, nor cytochrome C were detected in a media only condition (0) where no cells were plated (Fig. 1D, Supplementary Fig. S1).

**Figure 1.**
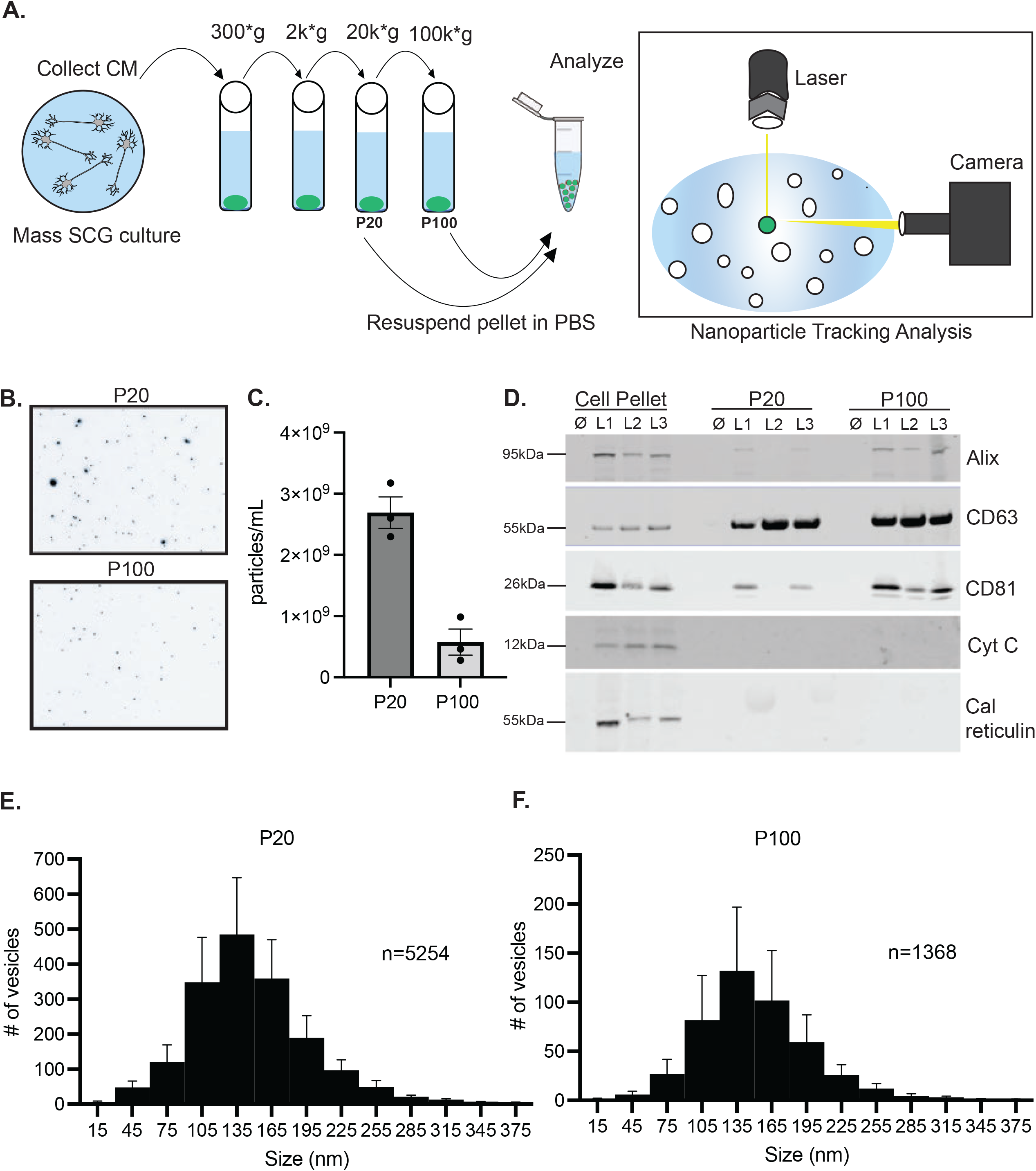
EV isolation and analysis by immunoblot and NTA. **A.** Schematic of EV isolation from SCG primary culture via ultracentrifugation and downstream nanoparticle tracking analysis (NTA) by ZetaView. Conditioned media is first spun at a low speed to remove contaminating cells. Next the supernatant is transferred to a fresh tube and spun at a higher speed. The supernatant is continually transferred and spun at increasing higher speeds. The pellets from the last two spins are collected and analyzed for EVs. **B.** Still frames captured from NTA ZetaView videos at t=30 secs. **C.** Quantification of the video analysis shown in B. Shown is mean ± SEM for 3 biological replicates measured at 11 positions, 3 cycles with two technical replicates. **D.** Immunoblot analysis of the canonical EV markers: Alix, CD63, CD81 and the intracellular markers: cytochrome C (mitochondria) and calreticulin (ER). Cell pellet, P20 and P100 fractions from three independent litters (L1, L2, L3) and a “no cell” media control (0) are shown. **E.** Size distribution histogram of 5,254 particles from the P20 fraction from 3 biological replicates. **F.** Size distribution histogram of 1,368 particles from the P100 fraction from 3 biological replicates.

To ensure the rigor of particle detection by NTA we conducted a series of solution controls including: 1. “complete media” (DMEM without phenol red, GlutaMAX, Prime XV IS-21, and 50ng/mL NGF), 2. 0.1μm filtered PBS (fraction resuspension solution), 3. Sterile 0.1μm filtered PBS incubated for 3 hours in centrifugation tubes, which are known to shed microplastics (UC tube = ultracentrifuge tube; MCT = microcentrifuge tube). NTA quantification of these control conditions showed that the concentration of particles derived from these sources is minimal (ranging from 1.16 × 10^6^ to 5.50 × 10^6^ particles/mL) (Supplementary Fig. S2). We also performed a “no cell” control to account for light scattering from microplastics shed by tissue culture plates. Complete media was added to the wells of an empty 12 well plate and left to incubate for 48 hours until it was processed by differential centrifugation and subjected to NTA (Supplementary Fig. S2). We observed 5.23 × 10^7^ ± 5.21 × 10^6^ particles per mL for the P20 fraction and 5.38 × 10^7^ ± 5.07 × 10^6^ particles per mL for the P100 fraction representing 1.94% and 9.36% of the particles observed in Figure 1C, respectively.

Next, we determined the optimal growth duration and minimum number of primary sympathetic cells necessary to robustly produce and detect EVs. To this end, we cultured cells from the SCG at densities ranging from 0 to 160,000 cells per well for 2 DIV or 7 DIV in a 12 well plate and EV secretion in a 48-hour window was determined (Supplementary Fig. S2). As expected, increased starting cell density increased the number of particles detected in media from 2 DIV and 7 DIV cultures (Supplementary Fig. S2). For this study we chose to use 100,000 cells grown for 7 DIV because we could reliably detect EV markers by immunoblot (Supplementary Fig. S2) and greater than 3×10^9^ particles per mL by NTA at starting cell densities above 80,000 (Supplementary Fig. S2). Lastly, we employ microfluidic devices (MFDs) in this study and wanted to account for microplastics that could be shed into the media. To test this, we added complete media to MFDs which contained no cells and pooled the media from 1, 2, 4 or 10 MFDs which showed 1.28 × 10^7^, 2.23 × 10^7^, 4.86 × 10^7^, and 8.32 × 10^7^ particles per mL, respectively (Supplementary Fig. S2). Particle counts from microplastics were detected by NTA, but the counts were very low even when using 10 MFDs (Supplementary Fig. S2). Importantly, the particles shed from MFDs were more than an order of magnitude lower than the SCG conditions indicating that particles shed from these MFDs are not contributing in a significant way to the total concentration of counted particles.

We next assessed the size and morphology of SCG culture derived EVs using cryo-transmission electron microscopy (Cryo-EM). We collected high magnification micrographs of the P20 and P100 fractions that show EVs delimited by a membrane bilayer (Fig. 2A). Sizes were determined by measuring the diameter through the largest part of the vesicle. The full-size distribution histogram of both the P20 (black bars) and P100 (gray bars) fractions are shown separately and together (Fig. 2B). For the P20 and P100 fractions the mean EV diameter was 146 nm and 153 nm, and the median EV diameter was 115 nm and 95 nm, respectively. The size distribution histogram appears to have two distinct peaks: a narrow peak around 45 nm followed by a broader flatter shoulder of larger sizes (Fig. 2B, both fractions). This smaller sized EV population was not effectively resolved by NTA-based sizing (compare Fig. 2B to Fig. 1E,F) and only apparent from cryo-EM sizing. Next we collected low magnification micrographs of both the P20 and P100 fractions and found that the P20 fractions contained large electron dense aggregates that were difficult to measure as discrete vesicles (Supplementary Fig. S3). These aggregates were absent in the P100 fraction, most likely because the 20,000 x g spin effectively depleted large aggregated material before preparing the P100 fraction. Only vesicles that could be individually measured were included in size distribution histograms (shown in Fig. 2B). We also analyzed micrographs from P20 and P100 fractions from “no cell” controls (complete media that had undergone differential centrifugation) and found that no vesicles or large aggregates were detected by cryo-EM indicating that the observed aggregates are indeed biological material derived from cultured cells (Supplementary Fig. S3). The P20 fraction exhibits contamination by large aggregates, therefore we mostly restricted our analysis to P100 fractions going forward.

**Figure 2.**
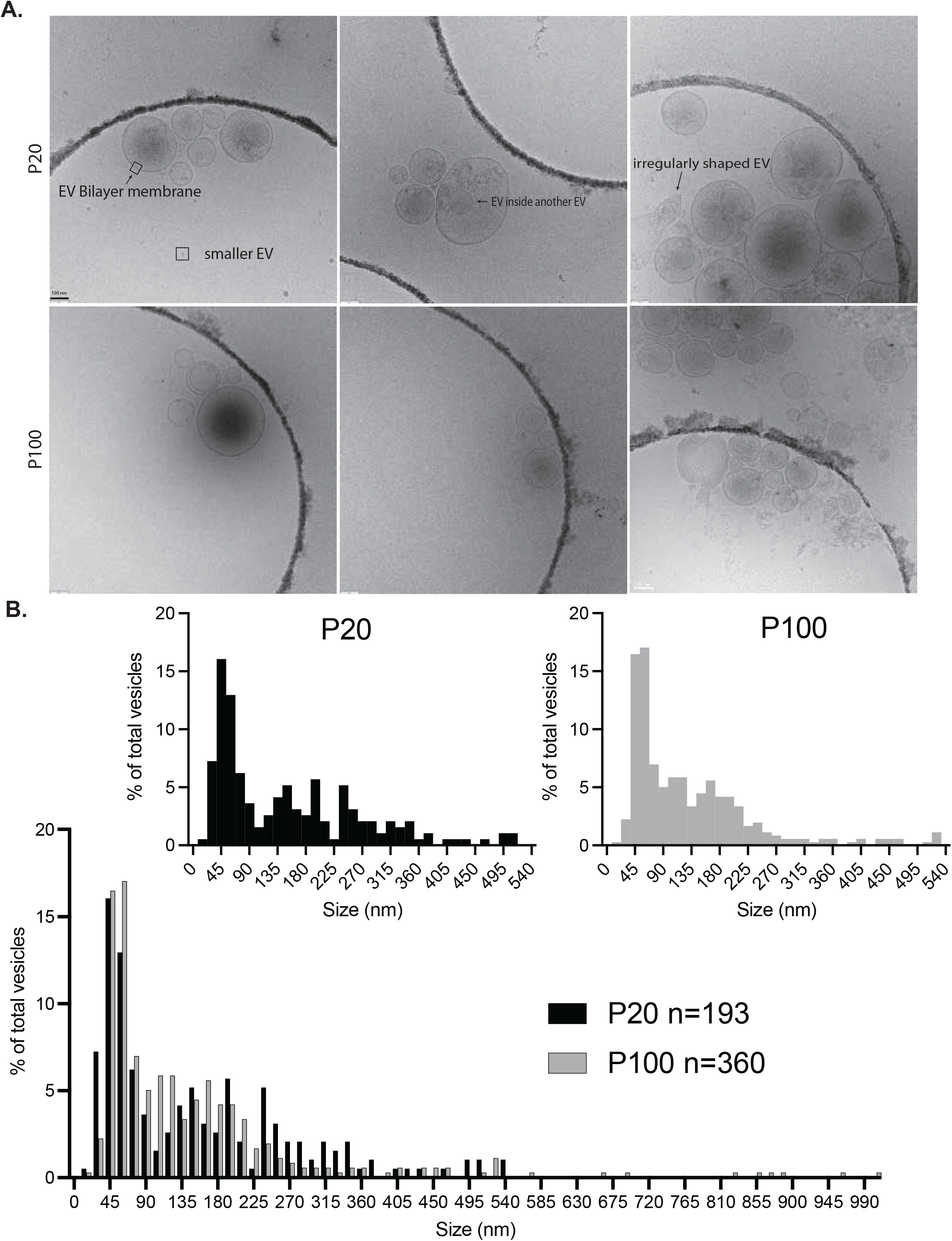
Morphology and Sizing of EVs by cryo-EM. **A.** Cryo-EM micrographs from the P20 (top row) and P100 (bottom row) fractions. Different types of EVs are annotated for easier appreciation of diverse EV morphologies. **B.** Size distribution histogram for all measured EVs from cryo-EM micrographs (P20: n=193, mean diameter 146.62 nm, n= 3 biological replicates; P100:n=360, mean diameter 152.59nm, n=3 biological replicates). Left histogram (black bars) is the sizing of the P20 fraction separated out from the combined histogram below. Right histogram (gray bars) is the sizing of the P100 fraction separated out from the histogram below. Scale bar is 100 nm for all images.

We next compared the size of the sympathetic EVs that we isolated with the size of those published in the literature. Since EVs can be derived from ILVs within MVBs after fusion with the plasma membrane, we decided to measure the size of ILVs within sympathetic MVBs from electron micrographs published in recent papers^11,17^. We found several micrographs in each paper containing sympathetic MVBs and measured their ILV sizes. ILV sizes ranged from 10 to 110 nm (Supplementary Fig. S4). Bronfman and colleagues showed micrographs of EVs derived from SCG neurons and NGF-differentiated PC12 cells^11^. Their EVs ranged from 30-100 nm in diameter (Supplementary Fig. S4). These published data align well with the first small peak visible in our data, consistent with the notion that these EVs are derived from MVB fusion and secretion of ILVs.

Further analysis of P20 and P100 micrographs revealed a vast heterogeneity in the morphology of EVs. We identified small vesicles that lacked a clear lipid bilayer (Supplementary Fig. S4) and termed them non-membranous vesicles (Supplementary Fig. S4). The EV field is increasingly reporting these small non-membranous EVs as exomeres or extracellular particles ^5,19,20^. Interestingly, the P20 fraction contained a larger percentage of these single membrane vesicles that were <60 nm in diameter compared to the P100 fraction (Supplementary Fig. S4). It is likely that these smaller particles are of higher density or aggregated with higher density particles and thus pellet in the P20 despite their small size. Additionally, EVs with diverse shapes and structures were detected with some EVs exhibiting long tubules (Supplementary Fig. S4) while other EVs were extremely electron dense (Supplementary Fig. S4). Lastly, several micrographs contained EVs that were inside of other EVs (Supplementary Fig. S4). There is speculation as to whether these EVs are naturally encapsulated inside each other or whether this is an artifact of ultracentrifugation resulting in membranes fusing into other membranes. However, this does not appear to be EVs imaged on a different z-plane from each other since their membranes curve or deform around other EVs (Supplementary Fig. S4). The size and number of EVs that were inside of other EVs are shown in Supplementary Fig. S4.

### Sympathetic EVs contain cargo derived from the distal axon

We next sought to determine if cargo originating in distal axons of SCG neurons could be recovered in EVs. To do this, we cultured SCG neurons in microfluidic devices (MFDs) which allowed us to separate the cell bodies (CB) of neurons from their distal axons (DA) by a series of microgrooves (Fig. 3A). Next, under fluidic isolation, we added an Alexa-conjugated wheat germ agglutinin (WGA), a well-known neuronal tracer, to the DA chamber to label SCG neurons at their distal axons (Fig. 3A). We collected conditioned media (CM) from the CB chamber and isolated EVs by differential centrifugation 15 hours after adding WGA-AF488 (Fig. 3A). The ZetaView NTA instrument is equipped with a filter allowing measurement of fluorescently labeled particles. We employed this to measure the total number of particles secreted (scatter) from the SCG neurons and the number of WGA^positive^ particles (fluorescent) (Fig. 3 B,C). By scatter, 4.05 × 10^9^ ± 9.71 × 10^8^ particles/mL were detected with 2.01 × 10^8^ ± 1.99 × 10^7^ WGA^positive^ particles/mL (Fig. 3C). Based on these findings we conclude that cargo originating in the distal axon can retrogradely traffic in the axon and be released as EVs from the somatodendritic domain.

**Figure 3.**
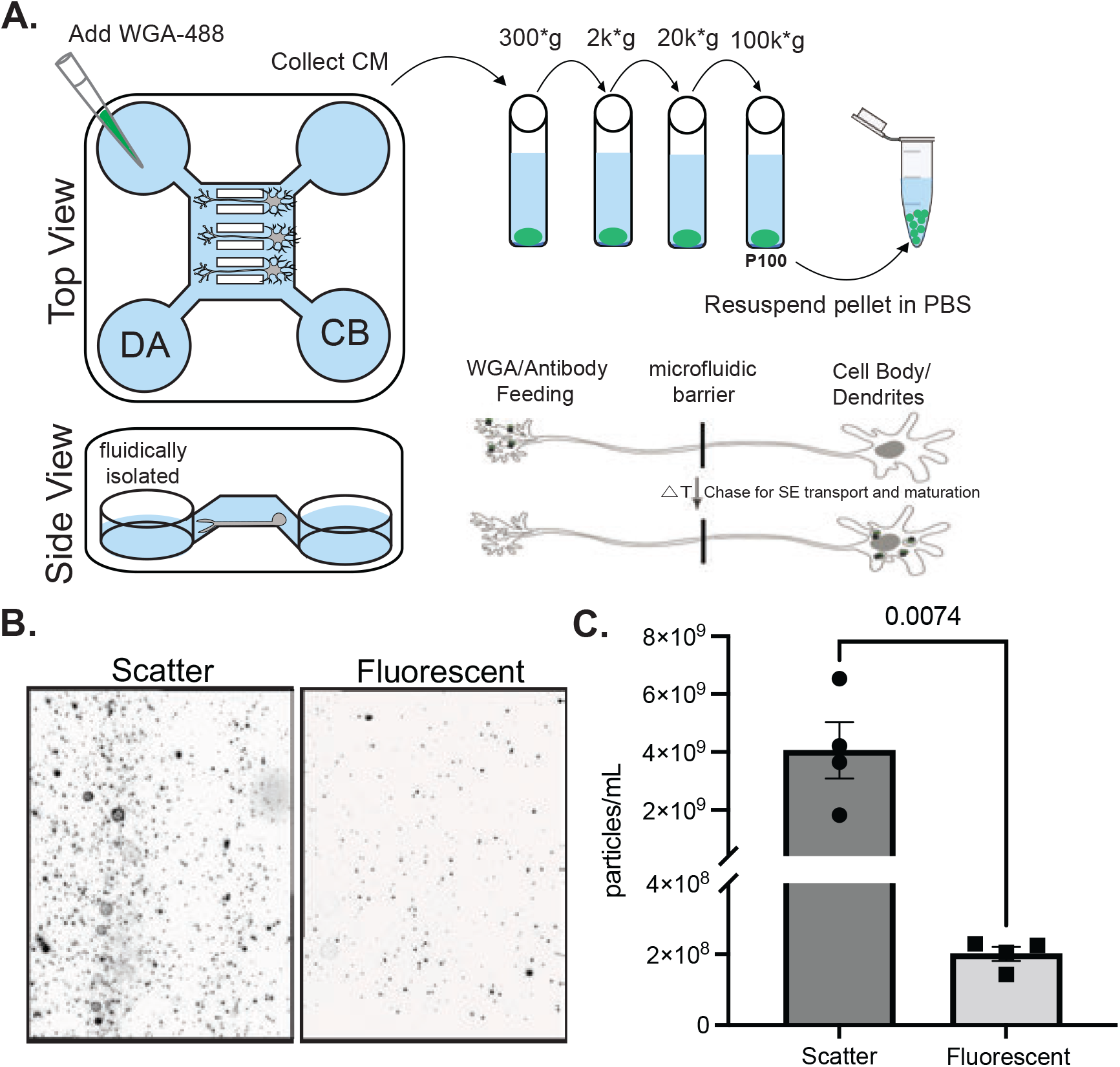
Somatodendritcally secreted sympathetic EVs carry cargo originating in the distal axon. **A.** Schematic of the WGA-488 feeding assay in microfluidic devices. WGA-488 was added to the distal axon (DA) chamber which was fluidically isolated from the cell body (CB) chamber. **B.** Still frames captured from NTA ZetaView videos at t=30 secs in scatter and fluorescent mode. **C.** Quantification of the total number of particles (scatter) and the number of fluorescent (WGA-488+) particles collected from the P100 fraction after WGA-488 addition to the DA chamber of MFDs containing wildtype SCG neurons. Shown is mean ± SEM for 4 biological replicates measured at 11 positions, 3 cycles with two technical replicates.

### Sympathetic EVs contain active retrogradely trafficked TrkA receptors

Once we determined that axonally derived cargos could be secreted in EVs, we speculated whether another retrogradely trafficked cargo, TrkA, could be secreted on EVs. TrkA bound to its high affinity ligand, NGF, is internalized into a signaling endosome (SE) in the axon terminals of developing SCG neurons ^21,22^. This SE is then retrogradely trafficked through the axon to the somata and dendrites ^23,24^. Here the SE can undergo endosomal maturation and be included in multivesicular bodies, the origin compartment for exosomes ^17^. We and others have also reported that retrograde SEs can fuse with the somatodendritic plasma membrane ^18,25^, which may release TrkA that might be present on ILVs. Thus, we wanted to determine if TrkA from the distal axons of sympathetic neurons could be recovered in EVs. Using immunoblotting of P20 and P100 EVs derived from SCG cells grown in mass culture (three separate litters L1-L3), we detect total TrkA (Fig. 4A) on purified EVs. Remarkably, TrkA is largely restricted to the P100 fraction suggesting that it is specifically packaged into smaller EVs (Fig. 4 A,). Next, we wanted to determine if TrkA secreted on EVs was activated, i.e. phosphorylated. We repeated the experiment with three additional litters and probed Western blots for phosphorylated TrkA (Fig. 4B). We observed that pTrkA was detectable in the P100 EV fraction, indicating that some activated TrkA is secreted on EVs.

**Figure 4.**
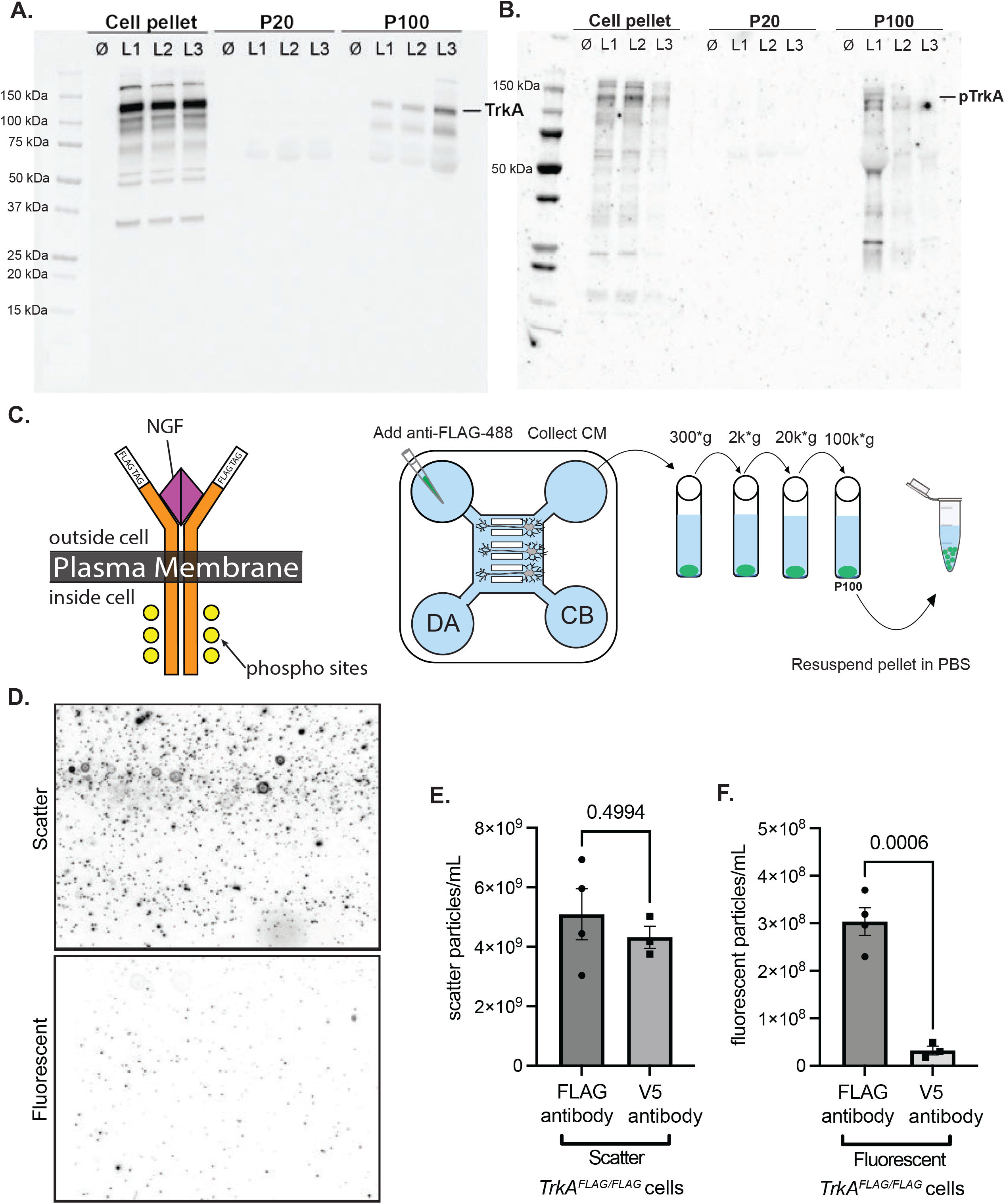
Somatodendritically secreted sympathetic EVs contain the TrkA receptor that originated in the distal axon and was transported retrogradely to the soma. **A.** Full immunoblot analysis of TrkA in cell pellets, P20 and P100 EV fractions from three independent litters (L1, L2, L3) and a “no cell” media control (0). **B.** Full immunoblot analysis of phosphorylated TrkA (Y-490) in cell pellets, P20 and P100 EV fractions from three independent litters (L1, L2, L3) and a “no cell” media control (0). Note: the three litters from **A.** are different from the three litters from **B. C.** Schematic showing the topology of FLAG-TrkA expressed from the mouse knockin locus in *TrkA^FLAG/FLAG^* mice (NGF: magenta triangle, phosphorylation sites: yellow circles). Sympathetic neurons from *TrkA^FLAG/FLAG^* mice were grown in compartmentalized microfluidic devices. Anti-FLAG antibody conjugated to AF488 was fed to the distal axons (DA) and 15 hours later conditioned media (CM) was collected from the cell body (CB) compartment and EVs were isolated by differential centrifugation. **D.** Still frames captured from NTA ZetaView videos at t=30 secs in scatter (top) and fluorescent (detecting Anti-FLAG-AF488) (bottom) mode. **E.** Quantification of the number of particles (scatter) collected from the P100 fraction after either Anti-FLAG-AF488 or V5-AF488 addition to the DA chamber of MFDs containing TrkA^FLAG/FLAG^ SCG neurons. Shown is mean ± SEM for 4 biological replicates measured at 11 positions, 3 cycles with two technical replicates. **F.** Quantification of the number of fluorescent (Anti FLAG-AF488+) particles collected from the P100 fraction after either Anti-FLAG-AF488 or V5-AF488 addition to the DA chamber of MFDs containing TrkA^FLAG/FLAG^ SCG neurons. Shown is mean ± SEM for 4 biological replicates measured at 11 positions, 3 cycles with two technical replicates.

In order to determine whether distally derived TrkA could be packaged into EVs upon retrograde arrival to the soma, we employed the MFD set up as described in Fig. 3A. We made use of a mouse line containing a FLAG tag that was knocked in frame with the extracellular domain of the TrkA locus (*TrkA^FLAG/FLAG^* transgenic mice) (Fig. 4C) ^26^. This allows selective labeling of pre-endocytotic TrkA receptors on the distal axon plasma membrane by adding an anti-FLAG antibody conjugated to AlexaFluor 488 just to the DA chamber of the MFD as previously reported (Fig. 4C)^27^. Using the paradigm set up in Figure 4C, we waited fifteen hours after anti-FLAG-AF488 antibody addition to the DA to collect the conditioned media from the CB chamber to isolate EVs (Fig. 4C). Using scatter and fluorescent NTA we detected 5.09 × 10^9^ ± 8.56 × 10^8^ total particles/mL and 3.03 × 10^8^ ± 2.91 × 10^7^ fluorescent particles/mL after the addition of anti-FLAG-AF488 antibody (Fig. 4D-F). To ensure that these fluorescent particles were indeed anti-FLAG antibody bound to TrkA and not due to non-specific uptake of anti-FLAG antibody, we added an irrelevant anti-epitope antibody, anti-V5-AF488, to *TrkA^FLAG/FLAG^* cells (Fig. 4E,F). As expected, the V5 antibody did not significantly change the total number of particles secreted (4.32 × 10^9^ ± 3.71 × 10^8^ particles/mL, Fig. 4E) nor did it result in significant fluorescent particle detection (3.24 × 10^7^, ± 9.16 × 10^6^ particles/mL, Fig. 4F) indicating that the anti-V5 antibody was not non-specifically endocytosed at significant levels. Next, we added anti-FLAG-AF488 antibody to wild type SCG cultures which do not express the *TrkA^FLAG/FLAG^* gene. We did not detect significant fluorescent particles thus confirming that the anti-FLAG-AF488 needed to bind the FLAG epitope on the FLAG tagged TrkA receptor to be internalized and secreted as EVs (Supplementary Fig. S5).

### Inhibition of phosphoinositide 3-kinase and phospholipase-C-γ reduces the number of TrkA EVs

The TrkA SE transduces its signal via several canonical downstream pathways such as phosphatidylinositol-3-kinase (PI3K) and phospholipase-C-γ (PLC-γ) (Fig. 5A). Interestingly, these downstream pathways often determine maturation and the trafficking fate of the SE ^28,29^. We next tested whether these pathways are important for TrkA^positive^ EV release. Using *TrkA^FLAG/FLAG^* SCG neurons cultured in MFDs, we simultaneously added anti-FLAG-AF488 to the DA chamber and different inhibitors to the CB chamber for 15 hours and then prepared EVs by differential ultracentrifugation from supernatant collected from the CB chamber (Fig. 5B). Inhibition of PI3K using the chemical inhibitor, LY294002, did not change the total number of particles secreted (Fig. 5C), but did cause a significant decrease in the number of fluorescent TrkA particles detected in the conditioned media (Fig. 5D) compared to a DMSO control. Interestingly, inhibition of PLC-γ using U73122 caused a significant increase in the total number of particles detected (Fig. 5E). Despite this increase in total particles/mL in U73122-treated MFDs, there was a significant decrease in the number of fluorescent particles compared to control (Fig. 5F). Immunocytological analysis of the cell bodies and dendrites of the neurons after the different drug treatments did not show any obvious morphological differences or cell loss that might account for the differences in EV secretion (Supplementary Fig. S6).

**Figure 5.**
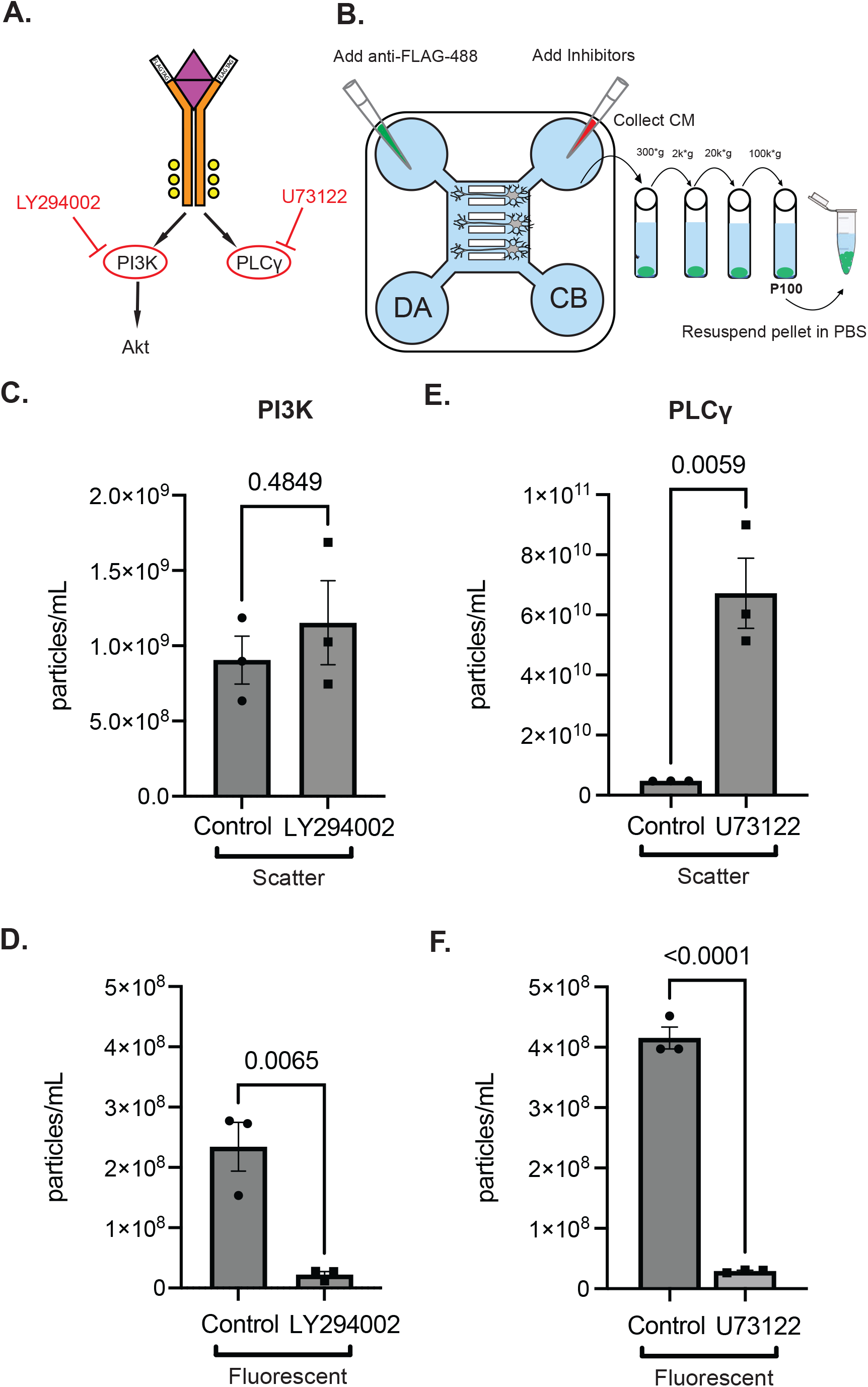
Inhibition of TrkA dependent signaling pathways suppresses TrkA^pos^ EV release. **A.** Schematic of the downstream pathways activated by TrkA phosphorylation and drugs that inhibit them. **B.** Schematic of drug and Anti-FLAG antibody application to compartmentalized neurons to generate TrkA^pos^ EVs. **C.** Quantification of the total number of particles (scatter) collected from the P100 fraction after anti-FLAG antibody addition to the DA chamber of MFDs containing *TrkA^FLAG/FLAG^* SCG neurons treated with the PI3K inhibitor, LY294002. **D.** Quantification of the number of fluorescent (AF488+) particles collected from the P100 fraction after anti-FLAG antibody addition to the DA chamber of MFDs containing *TrkA^FLAG/FLAG^* SCG neurons treated with the PI3K inhibitor, LY294002. **E.** Quantification of the total number of particles (scatter) collected from the P100 fraction after anti-FLAG antibody addition to the DA chamber of MFDs containing *TrkA^FLAG/FLAG^* SCG neurons treated with the PLC γ inhibitor, U73122. **F.** Quantification of the number of fluorescent (AF488+) particles collected from the P100 fraction after anti-FLAG antibody addition to the DA chamber of MFDs containing *TrkA^FLAG/FLAG^* SCG neurons treated with the PLCγ inhibitor, U73122. Shown is mean ± SEM for 3 biological replicates measured at 11 positions, 3 cycles with two technical replicates for panels **C-F**. P values are indicated above the pairwise brackets.

## Discussion

There are very few studies exploring EVs derived from the peripheral nervous system. Here we characterize EVs secreted from SCG cultures and conduct appropriate controls in accordance with the guidelines set forth by the ISEV ^12^. Using differential centrifugation/ultracentrifugation, we found that both the P20 and P100 fraction contain detectable amounts of canonical EV markers: CD63 and CD81, but not of the mitochondrial marker, Cytochrome C, or the ER marker calreticulin, suggesting that both these fractions are free from intracellular contamination (Fig. 1D). Another EV marker, Alix, is enriched in the P100 fraction relative to the P20. As expected, NTA revealed a higher concentration of EVs in the P20 fraction compared to the P100 fraction (the 100,000 x g pellet of the supernatant from the P20 fraction) (Fig. 1B,C). Low magnification cryo-electron micrographs revealed large aggregates of membrane material in the P20 fraction that are absent in the P100 fraction. Taken together these findings are consistent with previous work that the P20 fraction contains large dense EVs like apoptotic bodies or aggregates in addition to individual low density EVs like exosomes. The P100 fraction, in contrast, is depleted of large particles and aggregates and represents a purer fraction of smaller EVs^13,30^.

We use several controls, including a “no cell media only” control in all experiments to ensure that we are characterizing biologically derived EVs (Supplementary Fig. S2). Additionally, we manipulated the source of the EVs by varying cell density and found that cell density and EV concentration are positively correlated. We showed that the number of days in culture also affected the concentration of EVs secreted. This reflects the importance of allowing cultures to stabilize before collecting EVs, as EV secretion is heavily impacted by cellular state. Additionally, it highlights the importance of consistency in all parameters related to EV collection (DIV, density, duration of media conditioning) to accurately compare EV secretion across different conditions or genotypes. These parameters have been thoroughly described within the MISEV guidelines^12^.

Size analysis of sympathetic EVs by NTA and cryo-EM shows that the majority of EVs fall below 300 nm in diameter. The resolution limit of the ZetaView NTA is around 70-90 nm therefore sizing analysis excludes these smaller vesicles^31,32^. In contrast, cryo-EM detects the smaller EVs, but due to aggregation and concentration issues, larger EVs are excluded from analysis. Cryo-EM sizing shows two distinct peaks for both the P20 and P100 fraction, a sharper taller peak centered around 45 nm and a broader, wider peak around 180 nm (Fig 2B). We compared these findings with the size of published sympathetic EVs and found that EVs derived from NGF-differentiated PC12 cells and primary sympathetic cultures were below 100 nm in diameter^11^. Furthermore, we measured intraluminal vesicles in MVBs taken from micrographs of sympathetic neurons and found that the mean size was 52.8 nm and 79.9 nm ^11,17^. Based on these data, our findings align with the reported size of ILV-derived exosomes secreted by sympathetic neurons.

The origin and packaging of EV cargos in neurons is still relatively unknown ^3^. Endosomes are one of the intracellular compartments for trafficking EV cargos within the cell prior to secretion. Due to their high motility, endosomes can traffic cargo both locally and across long distances within neurons^33^. We used a fluorescently labeled lectin, wheat germ agglutinin (WGA), to visualize its trafficking through the cell from the distal axon to the cell body^34^. We show that WGA that originated in the distal axon can be packaged into EVs released at the cell body. WGA is primarily used as a neuronal circuit tracer to trace interconnected neurons;^35^ however, the mechanism by which WGA traverses synapses is still debated^36^. Our sympathetic neuronal cultures contain a mixture of all superior cervical ganglion cells types including satellite glia cells which outnumber the sympathetic neurons. Since we only added WGA to the DA of MFDs it is only being internalized by axons of neurons that grew through the microgrooves to reach the DA chamber. We show that a small proportion of all EVs secreted from our sympathetic cultures contain WGA and therefore are of neuronal origin. We speculate that these neuronal derived EVs might serve as one mechanism by which WGA can “hop” the synapse from one neuron to another. We note that we cannot determine what proportion of neuronally secreted EVs contain WGA since the CB chamber from which supernatant is collected contains many neurons whose axons did not traverse the microgrooves and were thus not able to pick up WGA. In addition, satellite glia growing in the CB chamber might also secrete EVs (see BLBP staining in Supplementary Figure S6). Future experimentation will be needed to more precisely determine the proportion of somatodendritically secreted EVs that contain retrograde cargo transported from the axon.

Retrograde trafficking of TrkA, is essential for sympathetic neuron survival^37,38^. The TrkA signaling endosome has recently been shown to arrive in the cell body as a multivesicular body; the precursor organelle for exosome formation^17^. In this study, newly arrived TrkA in the cell body after distal anti-FLAG antibody feeding was primarily found in MVBs (92.8%) at 1 hour with 80% found on ILVs^17^. For at least 8 hours, the majority of newly arrived TrkA remained associated with MVBs (>50%) with about 40% of the TrkA being found on ILVs^17^. Furthermore, these TrkA^positive^ MVBs were shown to contain activated TrkA receptors that remain in complex with PLC-ɣ^17^. Additionally, we have shown that the retrogradely trafficked TrkA SE, can undergo NGF-dependent recycling to the plasma membrane ^18^. Based on these data, we speculated that activated TrkA receptors could be packaged into EVs. Others have shown in sympathetic neurons, that another neurotrophin receptor, p75NTR, is routed away from the lysosome and towards secretion in EVs^11^.

Indeed, using compartmentalized cultures and *TrkA^FLAG/FLAG^* transgenic mice, we show that TrkA originating in the distal axon is retrogradely trafficked to the cell body in SEs and then subsequently secreted in EVs. While a potential function for TrkA^positive^ EVs is not known, we speculate that these retrogradely trafficked and secreted TrkA receptors are targeted to upstream recipient preganglionic neurons in the ganglia. A proportion of these TrkA-containing EVs remain catalytically active as evidenced through phosphorylated TrkA immunoblots possibly to aid in recipient neuron signaling. In addition, satellite glia found in close apposition to neuronal cell bodies in the SCG might also be recipients of TrkA^positive^ EVs. Future work is needed to more fully explore these possibilities.

Binding of NGF to TrkA triggers autophosphorylation of tyrosine residues located on its intracellular domain thereby initiating a kinase cascade on the downstream effector signaling pathways: Ras/MAPK, PI3K and PLC-ɣ^37,39–42^. Activation of different downstream pathways of TrkA have been shown to affect the trafficking and functional output of the TrkA SE. For example, blocking PI3K in the distal axon prevents the initiation of retrograde transport of the TrkA SE^29^. Additionally, recruitment of PLC-ɣ to the TrkA receptor promotes internalization of the ligand-receptor complex in the distal axon^28^. Based on these data, we wanted to test whether TrkA signaling affects the production of TrkA containing EVs. Our findings suggest that inhibition of PI3K signaling in the cell body of sympathetic neurons influences the production of TrkA^positive^ EVs without affecting total EV secretion. Additionally, inhibition of PLC-ɣ signaling in the cell body of sympathetic neurons reduces TrkA^positive^ EVs, but also causes a striking increase in the total number of EVs released. The mechanisms underlying these inhibitor’s effects on EV secretion remain unknown.

In summary, we have rigorously characterized EVs derived from primary sympathetic cultures through protein analysis, cryo-electron microscopy and nanoparticle tracking analysis. We have shown that EVs released from sympathetic cultures are heterogenous in size and morphology, and that our findings agree with and expand the sparse literature on sympathetic EVs. Finally, we demonstrate successful isolation of labeled EVs from specific neuronal domains. Specifically, we demonstrate that TrkA internalized at the distal axon can be secreted in EVs from the somatodendritic domain. Future studies investigating the mechanisms underlying TrkA partitioning into secretion competent organelles would help elucidate the effects different inhibitors have on TrkA EV numbers. Our results thus demonstrate a novel trafficking route for TrkA: it can travel long distances to the cell body, be packaged into EVs and secreted. Secretion of TrkA via EVs appears to be regulated by its own downstream effector cascades, raising intriguing future questions about novel functionalities associated with TrkA^positive^ EVs. Functional and recipient studies in the future will clarify the purpose of secreting these active TrkA-containing EVs and their functional output in their intended recipient cells.

## Materials and Methods

| REAGENT or RESOURCE | SOURCE | IDENTIFIER |
| --- | --- | --- |
| **Antibodies** |  |  |
| Rabbit Anti- Cytochrome C | Abcam | Cat # ab133504;<br>RRID:AB_2802115 |
| Rabbit Anti- CD63 | Abcam | Cat # ab217345;<br>RRID:AB_2754982 |
| Alexa Fluor 680 AffiniPure<br>Donkey anti-Rabbit IgG | Jackson ImmunoResearch | Cat # 711-625-152<br>RRID: AB_2340627 |
| Rabbit Anti- Calreticulin | Cell Signaling Technology | Cat # 12238S<br>RRID: AB_2688013 |
| Sheep Anti- Tyrosine<br>Hydroxylase | Milipore | Cat # AB1542<br>RRID: AB_90755 |
| Alexa Fluor 790 AffiniPure<br>Donkey anti-Rabbit IgG | Jackson ImmunoResearch | Cat # 711-655-152<br>RRID: AB_2340628 |
| Rabbit Anti- CD81 | Cell Signaling Technology | Cat # 10037S<br>RRID:AB_2714207 |
| Rabbit Anti – Alix | Cell Signaling Technology | Cat # 92880<br>RRID:AB_2800192 |
| Rabbit Anti-BLBP | Abcam | Cat # ab279649; RRID:AB |
| Chicken Anti-MAP2 | EnCor Biotechnology | Cat # CPCA-MAP2<br>RRID: AB_2138173 |
| Anti- FLAG DYKDDDDK tag<br>(D6W5B) Rabbit mAb Alexa<br>R 488 | Cell Signaling Technology | 15008S Lot 2 |
| Anti-FLAG DYKDDDDK tag<br>(D6W5B) Rabbit mAb Alexa<br>R 488 | Cell Signaling Technology | 15008S Lot 3 |
| **Biological samples** |  |  |
| NGF | In house, purified from mouse salivary glands |  |
| Chemicals, peptides, and recombinant proteins |  |  |
| Poly-D-Lysine | Sigma | Cat # P7886 |
| Prime XV IS-21 | Sigma | Cat # 91142 |
| Hyaluronidase | Sigma | Cat # H3884 |
| Collagenase | Worthington | Cat # LS004196 |
| Laminin | Invitrogen | Cat # 23017-01 |
| BSA | Sigma | Cat # A9647 |
| Polyacrylamide gels 4-12% | Genscript | Cat # M00654 |
| Trypsin | Sigma | Cat # T4799 |
| WGA-AF488 | Fisher Scientific | Cat # W11261 |
| PBS | Gibco | Cat # 14190-144 |
| Milk | Lab Scientific | Cat # M0841 |
| Beta mercaptoethanol | BioRad | Cat # 161-0716 |
| DMEM no phenol | Gibco | Cat # 31053-028 |
| GlutaMAX | Gibco | Cat # 35050-061 |
| FBS | R & D Systems | Cat # S11195H |
| LY294002 | Sigma | Cat # L9908 |
| U-73122 | Sigma | Cat # U6756 |
| **Experimental models: Organisms/strains** |  |  |
| C57 BL/6J mice | Jackson Laboratory |  |
| TrkA<sup>FLAG/FLAG</sup> | Gift from D. Ginty | Harvard |
| **Hardware, Software and algorithms** |  |  |
| Odyssey CLx | LI-COR |  |
| Optima TLX Ultracentrifuge | Beckman-Coulter |  |
| Trans-Blot Turbo Transfer | Bio Rad |  |
| Image J |  | https://imagej.nih.gov/ij/ |
| Tecnai F20 Twin Electron Microscope | FEI |  |
| Prism 9 | Graphpad | graphpad.com |
| ZetaView PMX-120 | Particle-metrix | particle-metrix.com |
| Illustrator | Adobe | adobe.com |
| **Other** |  |  |
| Microcentrifuge tubes | USAscientific | Cat # 1415-2500 |
| Tissue culture plates | Fisher Scientific | Cat # 150628 |
| Polycarbonate centrifuge tubes | Beckman | Cat # 343778 |
| Sylgard 184 Silicone elastomer kit | Krayden | Cat # DC2065622 |

### Animals

All animal use complied with the Association for Assessment of Laboratory Animals Care policies and was approved by the University of Virginia Animal Care and Use Committee protocol #3422 (Winckler lab) and protocol #3795 (Deppmann lab). All mice were maintained on a C57Bl/6J background and males and females were mixed in all experiments. Mouse lines used: C57BL/6J and TrkA^FLAG/FLAG 26^.

### Primary sympathetic neuronal cultures

Superior cervical ganglia were micro dissected from postnatal day 3 (P3) mouse pups as previously described, and kept in ice cold DMEM until enzymatic digestion^27^. Ganglia were transferred to an enzymatic solution containing 0.01 g/mL BSA, 0.4 mg/mL hyaluronidase and 4 mg/mL collagenase for 20 mins at 37°C. This solution was aspirated off and replaced with a 2.5% trypsin solution for 15 mins at 37°C. Cells were then washed in DMEM containing 10% FBS 3x and then subjected to trituration using a P1000 pipette and then a P200 pipette. Cells were then spun down at 300 x g and resuspended in complete media. A small 10 μL aliquot of cells was counted on a hemocytometer. Cells were plated at a density no less than 100,000 cells in a 12 well plate that had been precoated with poly-D-lysine and 1mg/mL laminin and washed 3x with sterile PBS. Cells were kept in an incubator at 37°C at 10% CO_2_ and media was changed every 48 hours.

### Compartmentalized WGA feeding assay

Sympathetic neurons were dissected as described above and dissociated neurons were plated in microfluidic devices (MFDs) as previously described^21,22^. To encourage axonal crossing of the microgrooves, neurons were exposed to 30 ng/mL NGF in the CB chamber and 80 ng/mL NGF in the DA chamber. At 6 DIV, 150 μL of complete media was added to the CB chamber and 100 μL of WGA-AF488 (1:200) in complete media was added to the DA chamber. Conditioned media was collected from the CB chamber 15 hours after the addition of WGA-AF488 and EVs were isolated.

### Compartmentalized Anti-FLAG antibody feeding assay

Sympathetic neurons from *TrkA^FLAG/FLAG^* animals were dissected as described above and dissociated neurons were plated in microfluidic devices (MFDs) as previously described^43,44^. To encourage axonal crossing of the microgrooves, neurons were exposed to 30 ng/mL of NGF in the CB chamber and 80 ng/mL NGF in the DA chamber. At 6 DIV, 150 μL of complete media was added to the CB chamber and 100 μl of anti-FLAG DYKDDDDK Tag (1:200) in complete media was added to the DA chamber. Conditioned media was collected from the CB chamber 15 hours after the addition of anti-FLAG tag AlexaFluor 488 conjugate and EVs were isolated. For drug treatments, 100 μL of complete media containing drug inhibitors (LY294002 50 μM, U-73122 1 μM, or 0.1% v/v DMSO) were added to the CB chamber and 150 μL of anti-FLAG-AF488 antibody (1:200) in complete media was added the DA chamber.

### EV isolation and differential centrifugation

Conditioned media was collected from cells after 48 hours and placed into 1.5 mL microcentrifuge tubes on ice. In experiments using mass cultures, conditioned media from one 12 well plate was used. In experiments using MFDs, conditioned media collected from the CB chamber of 6 MFDs were combined. The conditioned media was then centrifuged at 300 x g for 10 mins at 4°C to pellet the cells. The supernatant was transferred to a clean 1.5 mL microcentrifuge tube and centrifuged at 2,000 x g for 10 mins at 4°C to pellet dead cells. The supernatant was transferred to a clean 1.5 mL microcentrifuge tube and spun at 20,000 x g for 30 mins. The pellet from this step is the P20 fraction. The supernatant was transferred to polycarbonate tubes subjected to ultracentrifugation at 100,000 x g_max_ (rotor: TLA 120.2; k -factor: 42; 53,000 rpm) for 70 mins at 4°C. The pellet from this step is the P100 fraction.

### Nanoparticle Tracking Analysis

NTA was conducted using the ZetaView PMX 120 equipped with a 488 nm laser and a long wave pass filter (cutoff 500 nm) and CMOS camera. Samples were diluted to 1 mL in PBS prior to analysis. Each sample was measured at 11 different positions over 3 cycles ensuring a minimum number of 1000 traces were recorded. Samples were recorded at 25°C, pH 7.0 with a shutter speed and camera sensitivity of 75 at 30 frames per second. Automatically generated reports of particle counts were checked and any outliers were removed to calculate the final concentration.

### Western Blot

All samples were lysed directly in 1.2X Laemmli sample buffer containing 5% BME and boiled for 5 mins. Laemmli sample buffer recipe: 4% SDS (10% (w/v), 20% glycerol, 120 mM 1M Tris-Cl (pH 6.8) and 0.02% (w/v) bromophenol blue in water. Sympathetic cultures were washed with PBS and lysed directly on the plate with 200 mL of 1.2X Laemmli sample buffer. P20 and P100 fractions were lysed directly in micro/ultracentrifuge tubes with 30 mL of 1.2X Laemmli. The sample buffer was pipetted up and down 50 times along the walls of the tubes to collect the entire pellet. Samples were run on 4-12 % polyacrylamide gels with 7 mL of cell pellet fractions and 15 mL of P20 and P100 fractions loaded per well. Protein gels were transferred to nitrocellulose membranes using the Trans-blot turbo, blocked in 5% milk for 1 hour and incubated in primary antibody (Alix 1:1000, CD63 1:1000, CD81 1:1000, Cytochrome C 1:5000, Calreticulin 1:4000) diluted in 5% milk 0.1% TBST overnight at 4°C on a rocker. Membranes were then washed 3 x with 0.1% TBST and secondary antibodies (1:20,000) diluted in 0.1% TBST were incubated for 1 hour at room temperature. Blots were imaged using the Odyssey CLx imager.

### Electron Cryo-Microscopy

Cryo-TEM was performed by the molecular electron microscopy core at UVA. P20 and P100 fractions were resuspended in 30 mL PBS. An aliquot of sample (~3.5 μL) was applied to a glow-discharged, perforated carbon-coated grid (2/1-3C C-Flat; Protochips, Raleigh, NC), manually blotted with filter paper, and rapidly plunged into liquid ethane. The grids were stored in liquid nitrogen, then transferred to a Gatan 626 cryo-specimen holder (Gatan, Warrrendale, PA) and maintained at ~180°C. Low-dose images were collected on a Tecnai F20 Twin transmission electron microscope (FEI {now ThermoFisher Scientific}, Hillsboro, OR) operating at 120 kV. The digital micrographs were recorded on a TVIPS XF416 camera (Teitz, Germany).

### Immunocytochemistry

Cells were fixed in 4% PFA for 20 minutes at room temperature in the MFDs. Cells were washed 3 times with 1X PBS and then blocked and permeabilized in 5% normal donkey serum and 0.2% TritonX-100 for 20 minutes. Primary antibodies were diluted in 1% BSA and applied overnight at 4 °C. Secondary antibodies were diluted in 1% BSA and added for 30 mins at room temperature. MFDs were washed 3x with 1x PBS and imaged on an inverted Zeiss 980 microscope with an Airyscan detector using a 40X oil objective (NA 1.3)

### Statistics and Measurements

Vesicles were measured at their widest diameter using the segment tool in Image J. Only vesicles that could be individually measured were included in size distribution histograms (shown in Supplementary Fig. S3). Statistical analyses were performed using Prism 9 software. All values are shown as mean ± SEM. Differences between samples were determined using unpaired, two-tailed t-tests. Statistical significance is a p value < 0.05. The p values are denoted on top of each bracket pair.

## Acknowledgements

Transmission electron micrographs were recorded at the University of Virginia Molecular Electron Microscopy Core facility (RRID:SCR_019031), which is supported in part by the School of Medicine and built with NIH grant G20-RR31199. We acknowledge the Keck Center for Cellular Imaging for the use of the Zeiss 880/980 multi-photon Airyscan microscopy system (PI-AP; NIH-OD025156). This work was supported by R21NS111991 awarded to CD and BW, 2T32GM007055-44 and F31NS126020 awarded to A.M and F32NS103770-02 awarded to A.K.

## Author Contributions

Study design and concept: A.M. A.K, B.W, and C.D. Data collection: A.M, A.K. Data Analysis: A.M., A.K., F.K. Data Interpretation: A.M., A.K, B.W., C.D. Manuscript and writing: A.M., B.W., C,D.

## Figure Legends

**Supplementary Figure S1.**
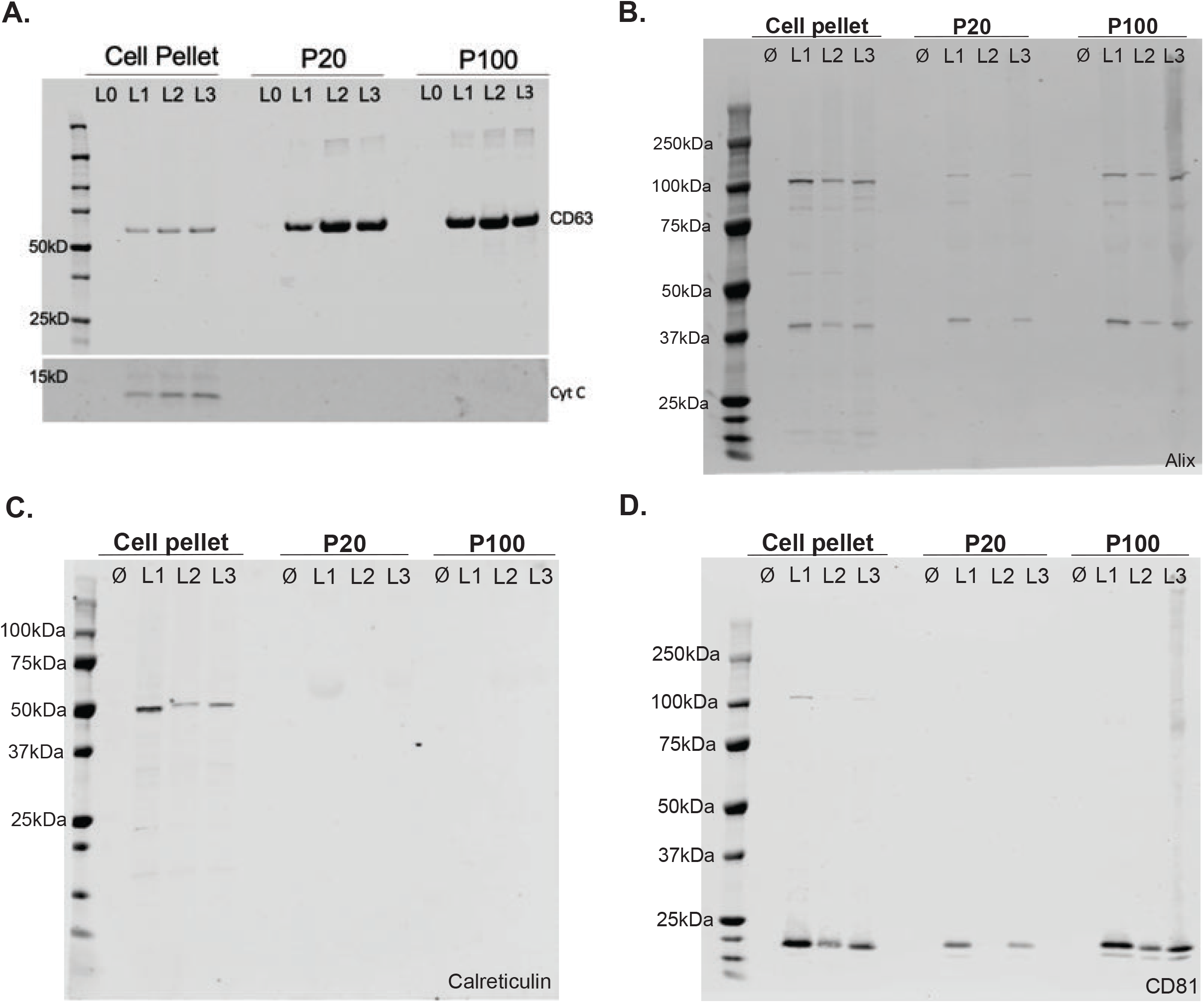
Immunoblots of EV and cell lysate markers shown in Fig. 1D. **A.** Full immunoblot that was physically cut in half before addition of primary antibody against the canonical EV marker tetraspanin, CD63 and the mitochondrial marker, Cytochrome C (Cyt C). **B.** Full immunoblot of the canonical EV marker and accessory ESCRT protein, Alix. Full length Alix is predicted to run around 95 kDa. The smaller band is either non-specific or represents a breakdown product of full length Alix. **C.** Full immunoblot of the endoplasmic reticulum marker, Calreticulin, only appearing in the lysate and not the EV fractions. **D.** Full immunoblot of the canonical EV marker tetraspanin, CD81 from the same blot in panel **B** that was stripped and re-probed.

**Supplementary Figure S2.**
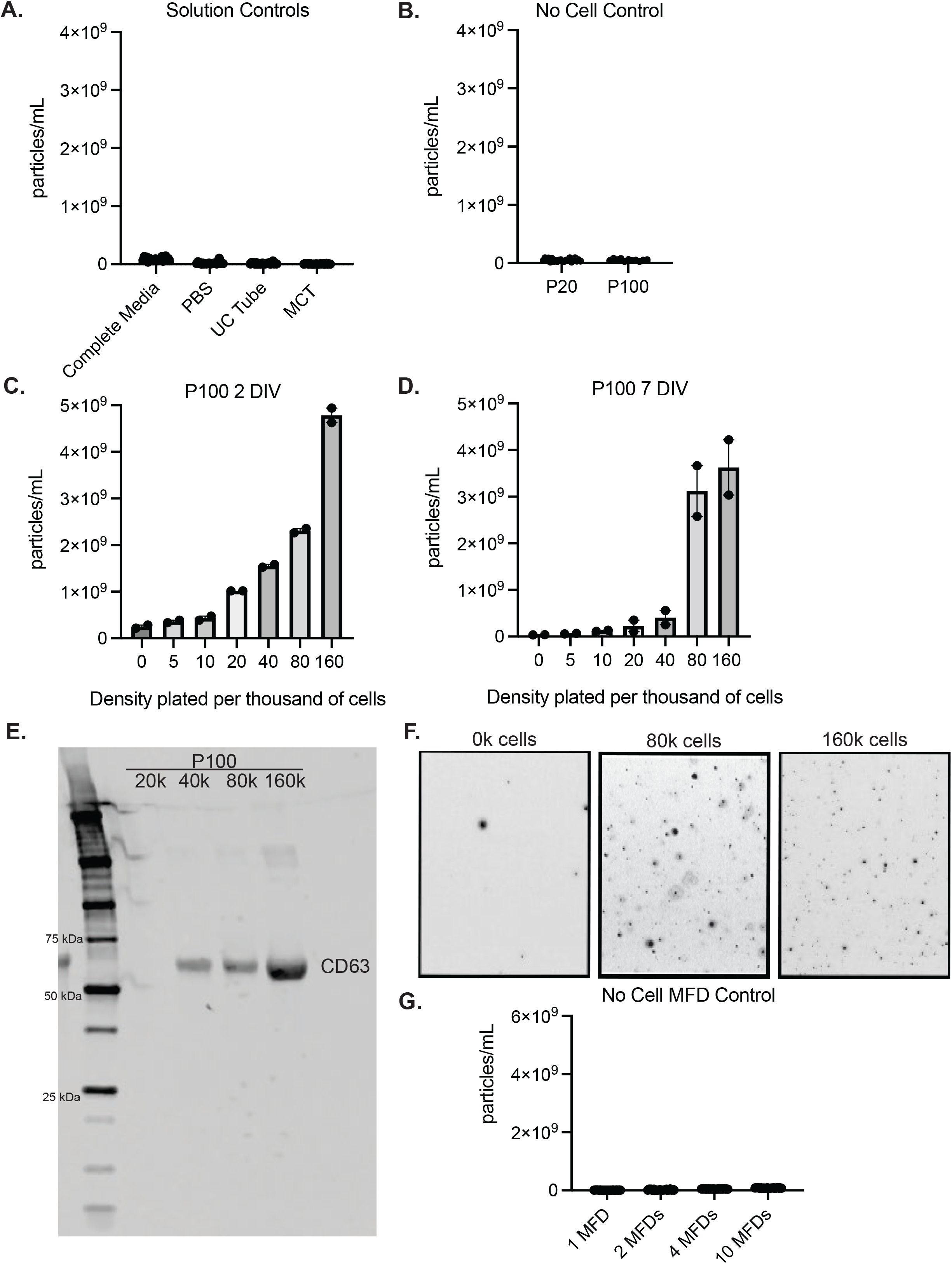
Density and days in vitro affect EV production. **A.** One milliliter of each undiluted solution condition was analyzed by ZetaView for non-EV scattering particles. Complete media (DMEM no phenol red, GlutaMAX, Prime XV IS-21, 50 ng/mL NGF), PBS (PBS), UC tube (PBS that sat in a polycarbonate centrifuge tube for 3 hours), MCT (PBS that sat in a microcentrifuge tube for 3 hours). Shown is mean ± SD for two technical replicates, 11 positions, 3 cycles. **B**. “No cell” only control consisting of complete media (DMEM no phenol red, GlutaMAX, Prime XV IS-21, 50 ng/mL NGF) that was plated in a 12 well plate and changed every 48 hours before collection and differential centrifugation. Shown is mean ± SEM for 3 biological replicates measured at 11 positions, 3 cycles with two technical replicates. **C.** Density and 2 DIV curve from P100 fraction. Cells were plated at the density shown on the x-axis and grown for 2 DIV before CM was collected for EV isolation and NTA analysis. Shown is mean ± SEM for 2 biological replicates measured at 11 positions, 3 cycles, two technical replicates. **D.** Density and 7 DIV curve from P100 fraction. Cells were plated at the density shown on the x-axis and grown for 7 DIV with media changes every 48 hours before CM was collected for EV isolation and NTA analysis. Shown is mean ± SEM for 2 biological replicates measured at 11 positions, 3 cycles, two technical replicates. **E.** Immunoblot analysis of CD63 at different densities of plated SCG cells. n=1 biological replicate. **F.** Still frames captured from NTA ZetaView videos at t=30 secs. EVs are from cells plated at 160,000 cells per well or 0 cells per well in a 12 well plate. **G**. “No cell” MFD control consists of complete media (DMEM no phenol red, GlutaMAX, Prime XV IS-21, 50 ng/mL NGF) that was plated and pooled from either 1, 2, 4, or 10 MFDs and changed every 48 hours before collection and differential centrifugation. Shown is mean ± SD for two technical replicates, 11 positions, 3 cycles.

**Supplementary Figure S3.**
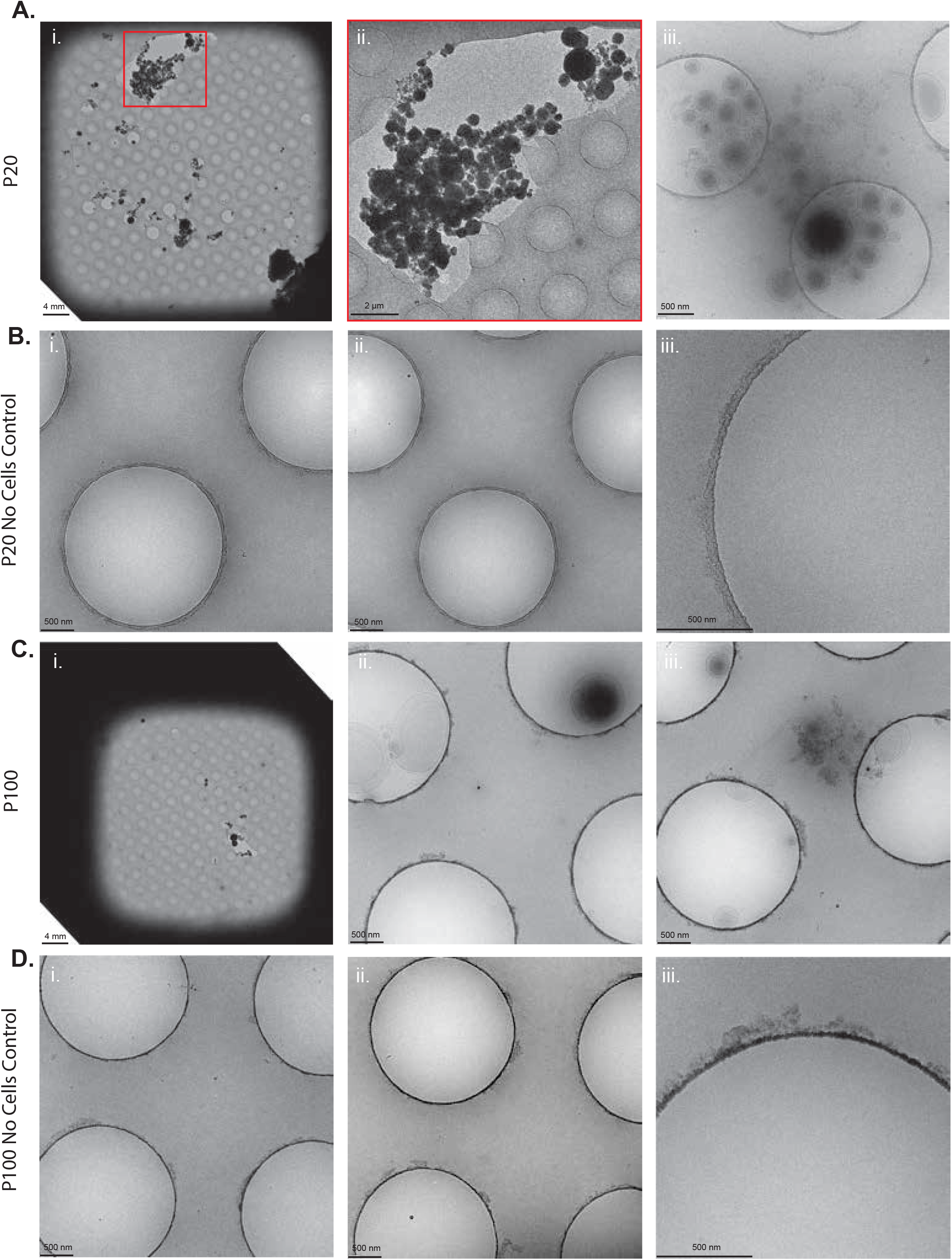
Low magnification cryo-EM micrographs of EVs. **A.** Low magnification micrographs of the P20 fraction. i. Shown are large aggregates which cannot be measured as discrete EVs. Scale bar is 4 mm. ii. Zoomed in view of the red boxed inset in i. Scale bar is 2 μm. iii. Discrete double membrane enclosed EVs are discernable with different sized EVs with different electron densities. Scale bar is 500 nm. **B.** Low magnification micrographs of the P20 “no cell” control fraction. i., ii., and iii. all show that no EVs are pelleted down from media that was added to tissue cultures dishes in which no cells were present. Scale bar is 500 nm for all. **C.** Low magnification micrographs of P100 fraction. i. Full grid view of P100 fraction with noticeably fewer large aggregates as compared to the P20 fraction. Scale bar is 4 mm. ii. EVs with interesting shapes and electron densities are viewable in the perforations. Scale bar is 500 nm. iii. Cluster of heterogeneous EVs. Scale bar is 500 nm. **D.** Low magnification micrographs of P100 “no cell” controls. (i., ii., iii.) all show that EVs are not sedimented after ultracentrifugation when conditioned media is collected from wells in which no cells were present. Scale bar is 500 nm for all.

**Supplementary Figure S4.**
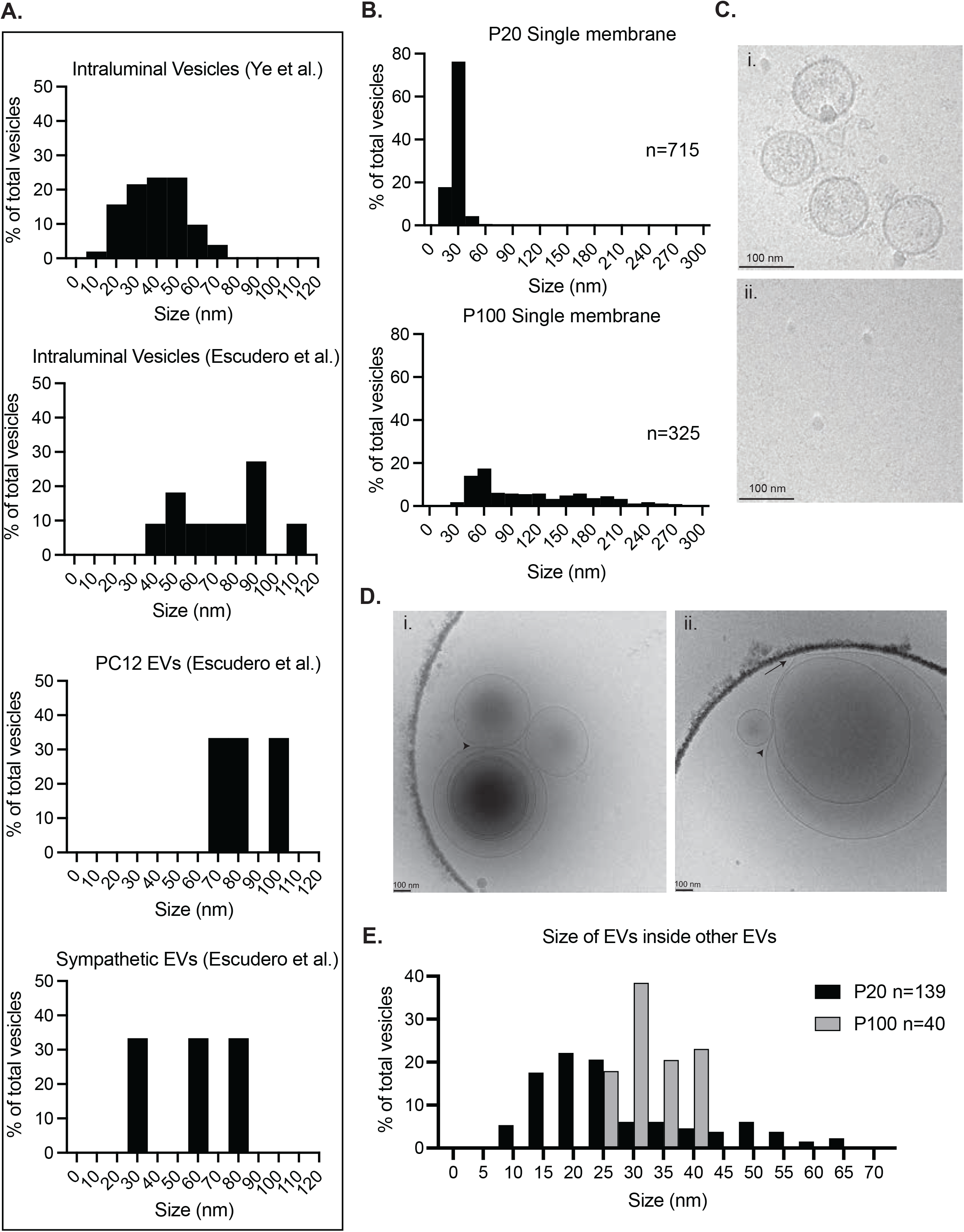
Heterogeneity in size and morphology of sympathetic EVs. **A.** Size distribution histogram comparing intraluminal vesicles (ILVs) from sympathetic neuron micrographs from Ye et al., 2018 ^17^ (mean diameter 52.8 nm) and Escudero et al., 2014 ^11^ (mean diameter 79.9 nm) as well as EVs derived from NGF-differentiated PC12 cells (mean diameter 80.0 nm) and sympathetic EVs from Escudero et al., 2014 ^11^ (mean diameter 58.3 nm). **B.** Size distribution histogram of single membrane enclosed EVs from the P20 fraction (top) (n= 715 EVs, mean ± SEM is 21.04 nm ± 15.59 nm, n=3 biological replicates) and P100 fraction (bottom) (n= 325 EVs, mean ± SEM is 159.03 nm ± 191.56 nm, n=3 biological replicates). **C.** Zoomed in micrograph of small EVs. i. Small EVs with a distinct double membrane lipid bilayer. Scale bar is 100 nm. ii. Sub 30 nm diameter exomeres with only a single membrane. Scale bar is 100 nm. **D.** Heterogeneity in size and structure of EVs. i. micrograph demonstrating EVs inside EVs (data quantified in E), electron dense EVs and EVs deforming around each other (arrowhead). ii. Micrograph showing EVs inside EVs, EV membranes deforming around each other (arrowhead) and long tubule-like projections from EV membranes (arrow). Scale bar is 100 nm for all images. **E.** Size distribution histogram of EVs enclosed inside of other EVs for both the P20 (mean ± SEM is 35.5 nm ± 26.79 nm) and P100 fraction (mean ± SEM is 24.59 nm ± 5.44 nm).

**Supplementary Figure S5.**
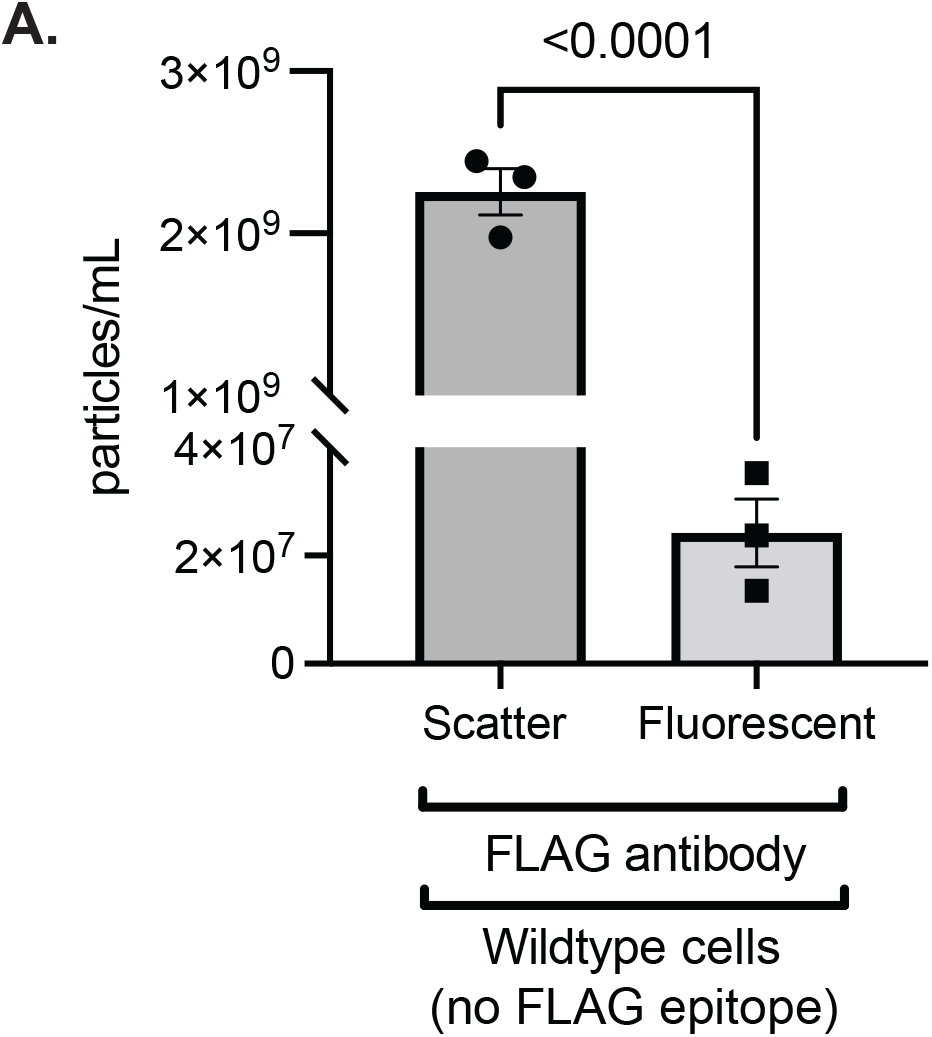
Controls for compartmentalized FLAG feeding assays. **A.** Quantification of the total number of particles (scatter) and the number of fluorescent (AF488+) particles collected from the P100 fraction after Anti-FLAG-AF488 addition to the DA chamber of MFDs containing wildtype (no FLAG epitope) SCG neurons. Shown is mean ± SEM for 3 biological replicates measured at 11 positions, 3 cycles with two technical replicates.

**Supplementary Figure S6.**
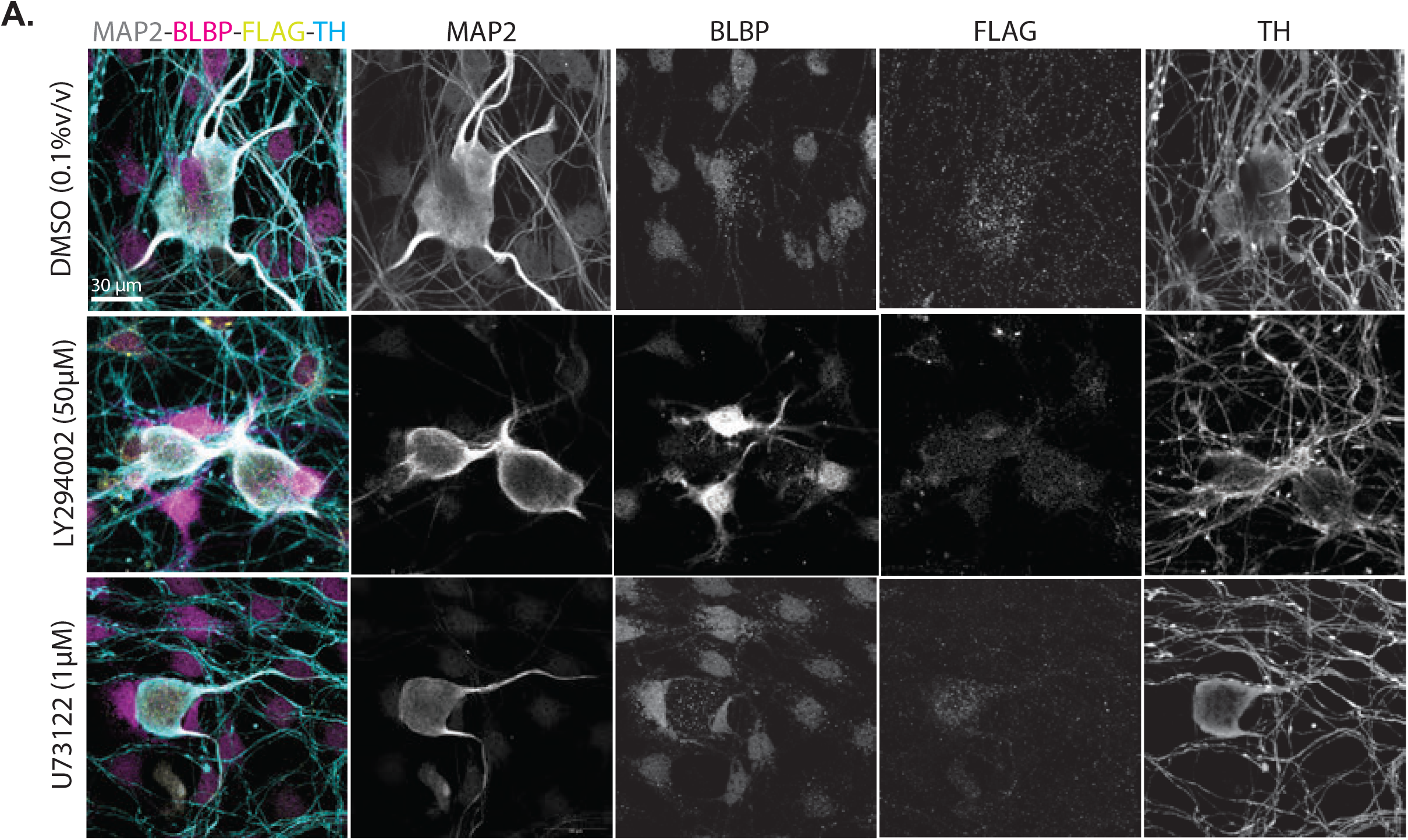
TrkA inhibitors do not affect SCG neuron morphology. A. Images of the cell bodies of compartmentalized SCG neurons treated with inhibitors in the cell body chamber and anti-FLAG-AF488 antibody in the distal axon chamber for 15 hours. Staining against MAP2 (somatodendritic domain), BLBP (satellite glia), anti-FLAG-AF488, and TH (SCG neuron). Scale bar is 30 μm.

